# Subjective perceptual experience tracks the neural signature of sensory evidence accumulation during decision formation

**DOI:** 10.1101/373464

**Authors:** Chiara F. Tagliabue, Domenica Veniero, Christopher S. Y. Benwell, Roberto Cecere, Silvia Savazzi, Gregor Thut

**Author notes:** These authors contributed equally to this study (Chiara F. Tagliabue, Domenica Veniero). **Corresponding author:** Chiara F. Tagliabue, CIMeC – Center for Mind/Brain Sciences, University of Trento, Corso Bettini 31, 38068, Rovereto, Italy.

## Abstract

How neural representations of low-level visual information are accessed by higher-order processes to inform decisions and give rise to conscious experience is a longstanding question. Research on perceptual decision making has revealed a late event-related EEG potential (the Centro-Parietal Positivity, CPP) to be a correlate of the accumulation of sensory evidence. We tested to what extent this evidence accumulation signal is driven by externally presented (physical) versus internally experienced (subjective) sensory evidence. The results show that the known relationship between external evidence and the evidence accumulation signal (reflected in the CPP amplitude) is mediated by the level of subjective awareness. Additionally, the CPP closely tracks the subjective perceptual evidence during both correct and incorrect trials. Hence, a remarkably close relationship exists between the evidence accumulation process (i.e. CPP) and subjective perceptual experience, suggesting that neural decision processes and components of conscious experience are tightly linked.

## Introduction

A central question in the study of decision making is how lower-level sensory information is accessed by higher-order processes to inform decisions and form conscious percepts. Research on perceptual decision making in human participants has established a late EEG potential characterized by a centro-parietal positivity (called CPP) as a neural correlate of sensory evidence accumulation [1, 2, 3]. The CPP is commensurate with the P300 family of event-related potentials (ERP) ([4]; for review, see [5]). In line with sequential information sampling models of decision making that assume integration of noisy evidence over time [6], the CPP build-up rate increases proportionally with the strength of the exogenously presented sensory evidence to then peak at the time when a decision has been reached [1]. The CPP thus shows the defining “build-to-threshold” features of a decision variable [7] and because it is neither specific to any particular sensory modality nor feature [1], nor to any motor requirements [2, 8, 9], it is believed to reflect evidence accumulation at an intermediate, abstract level of processing.

The process of evidence accumulation from early sensory stimulus representations depends not only on the strength of externally presented sensory evidence (stimulus intensity), but also on ‘internal’ sources of variability such as neural noise, in particular when the stimulus is weak. This can explain misperceptions (such as false alarms) in terms of erroneous evidence accumulation in sequential sampling models. Accordingly, it has been shown that the CPP is present not only for correct but also erroneous decisions [1, 8, 10]. Hence, the CPP tracks not only external evidence but also an internal decision quantity. In line with this, a growing number of studies have shown the CPP to reflect subjective aspects of the decision process such as decision confidence [11, 12, 13]. Interestingly, an evoked potential of similar latency and topography (Late Positivity, LP) has been shown to scale with perceptual awareness [14], hence to be associated with subjectively reported (experienced) evidence. Here, we wanted to directly compare to what extent the neural correlate of evidence accumulation, reflected in the CPP, is driven by externally presented (physical) versus subjectively experienced evidence. We therefore examined the CPP as to its co-variation with physical versus subjectively experienced stimulus intensity. We manipulated the amount of available sensory evidence and asked our participants to explicitly rate the strength of their subjective experience, here defined as the clarity of the percept (using a four point scale, the Perceptual Awareness Scale (PAS) [15]). Our data show that the CPP amplitude far more closely tracks the subjective reports of stimulus clarity than objective stimulus intensity, suggesting that a close relationship exists between the neural correlates of evidence accumulation and conscious awareness.

## Results

While recording EEG, we presented difficult-to-detect visual stimuli that were either brighter or darker than the background at three intensity levels (i.e. contrast-from-background), collected discrimination performance and then asked the participants to rate the clarity of their perception on the PAS scale (Fig 1.). This allowed us to investigate to what extent the CPP scales with the amount of external sensory evidence versus the level of subjective clarity of the percept (the internally experienced evidence), when the alternative variable is accounted for.

**Figure 1.**
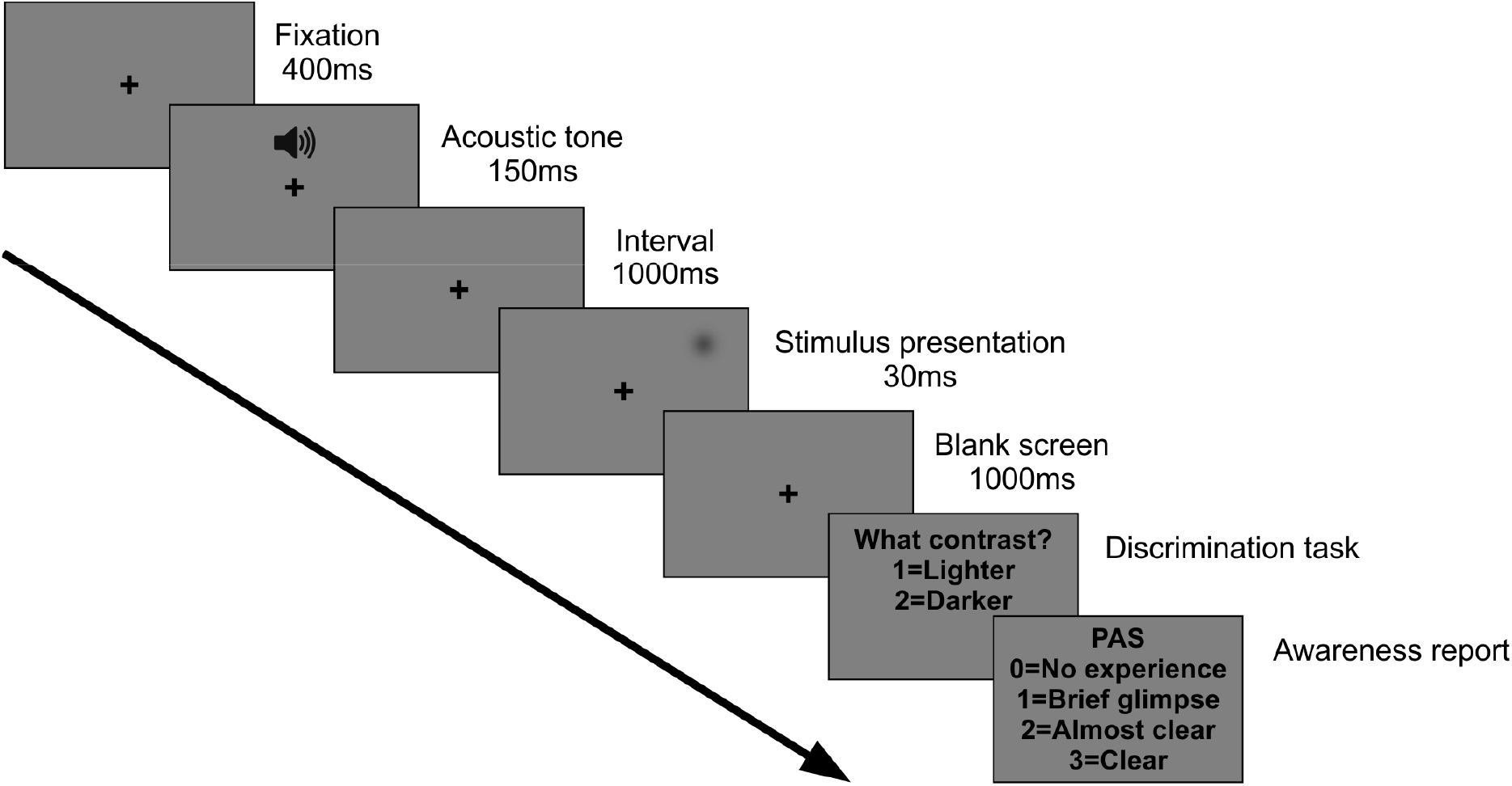
Single trial structure: Following an acoustic alerting tone, a brief visual stimulus was presented always at the same position in the upper right visual field. Stimuli could be either brighter or darker than the background and were presented at 3 different individually adjusted stimulus intensities (low, intermediate and high). After 1000 ms, participants were asked to report the brightness of the stimulus relative to the background (Discrimination task) and then rate the clarity of their perception on the Perceptual Awareness Scale (PAS) (Awareness report).

### Behavioral responses to visual stimuli

To manipulate the amount of sensory evidence, we varied stimulus intensity (low, intermediate and high) by selecting 3 different contrast levels (corresponding to stimuli presented at 25%, 50% and 75% of individual detection thresholds as determined prior to the experiment, see experimental procedures). Participants were asked to indicate the brightness of the stimulus relative to the background (“lighter” or “darker”, prompted by a first question screen) for assessing discrimination accuracy, and to then rate the clarity of their percept (“no experience”, “brief glimpse”, “almost clear” or “clear”, prompted by the second question screen) (Fig 1).

Participants performed the task reliably as illustrated by the distribution of subjective perceptual awareness ratings across the three intensity levels (Fig. 2a). For low intensity stimuli (orange line), participants indicated most often “no experience” (PAS=0), followed by a “brief glimpse” (PAS=1), “almost clear” experience (PAS=2), and very few “clear” experience (PAS=3). For intermediate intensity stimuli (purple line), the percentage of responses was more equally distributed across awareness rating levels. For high intensity stimuli (cyan line), participants indicated least often having “no experience” (PAS=0), followed by more frequent “brief glimpses” (PAS=1), “almost clear” (PAS=2) and “clear” experiences (PAS=3). Finally, catch trials were rated in 88.1% of trials as PAS=0 (“no experience”).

**Figure 2.**
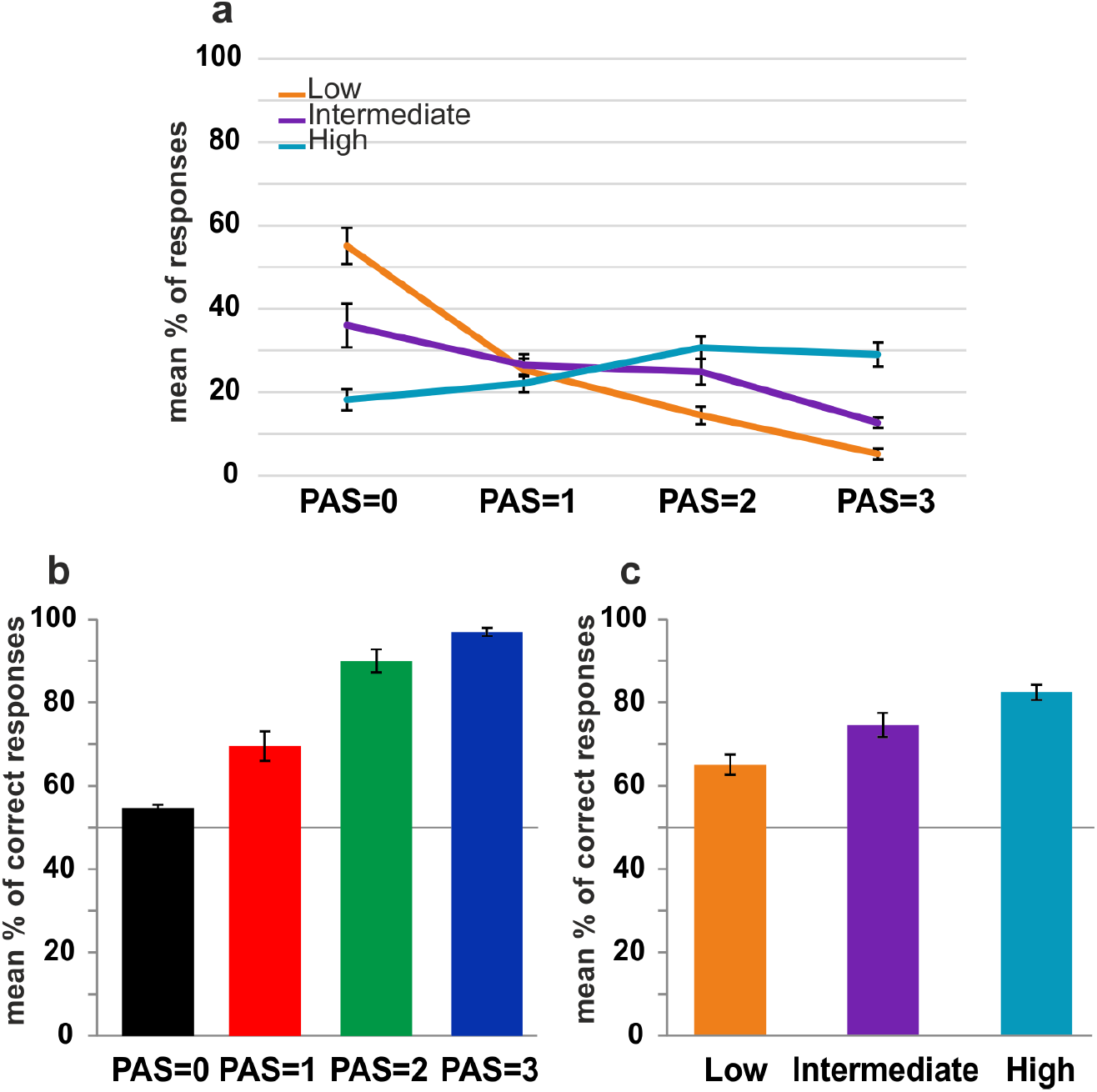
Behavioral results. **(a)** PAS rating variability for each level of external sensory evidence. Error bars represent standard errors. Mean percentage of correct discrimination responses as a function of **(b)** PAS ratings and **(c)** stimulus intensity. Error bars represent standard errors and the solid line (50%) chance level.

Sorting trials according to the clarity of subjective experience (i.e. PAS=0, PAS=1, PAS=2, PAS=3) revealed that as the clarity of the percept increased, accuracy also increased (Fig. 2b), as expected [repeated-measures ANOVA (degrees of freedom corrected using Greenhouse-Geisser estimates of sphericity): F(1.688,16.882) = 113.2, p < 0.01; linear trend F(1,10) = 1000.7, p < 0.01]. Similarly, sorting trials according to the strength of the presented evidence (i.e. low, intermediate, high stimulus intensity) showed that as objective sensory information increased, accuracy also increased (Fig. 2c) [repeated-measures ANOVA: F(2,20) = 35.5, p < 0.01; linear trend F(1,10) = 89.9, p < 0.01].

Overall, these results thus show that both factors of interest (visual awareness and stimulus intensity level), i.e. the internally experienced and externally presented sensory evidence, co-vary, but also show considerable trial-by-trial variability. Next, we examined to what extent the CPP amplitude is varying with each measure. In addition, we ran a mediation analysis to test whether the known relationship between stimulus intensity and CPP [1, 2, 3, 16] is mediated by subjective experience.

### Event-related potentials

We first plotted the CPP as a function of external physical evidence (stimulus intensity), not taking into account internally experienced evidence (PAS ratings) (Fig. 3). As expected, the CPP scaled with stimulus intensity. However, note the large variability around the mean per stimulus intensity (Fig. 3, shaded areas). To test whether this variability is explained by the variability in PAS ratings observed for each intensity level (see Fig. 2a), we identified the relative contribution of externally presented versus internally experienced evidence to the CPP. To this end, we tested whether the CPP amplitude varies with subjective awareness ratings when physical stimulus properties were held constant, or vice versa, whether this potential varies with physical stimulus properties when subjective ratings were held constant. That is, we compared ERP amplitudes evoked by different levels of subjective or objective evidence while controlling for the contribution of the alternative variable. To numerically equate the value of the alternative variable across all levels of comparison, we used random trial sub-sampling (see Supplementary Fig. 1).

**Figure 3.**
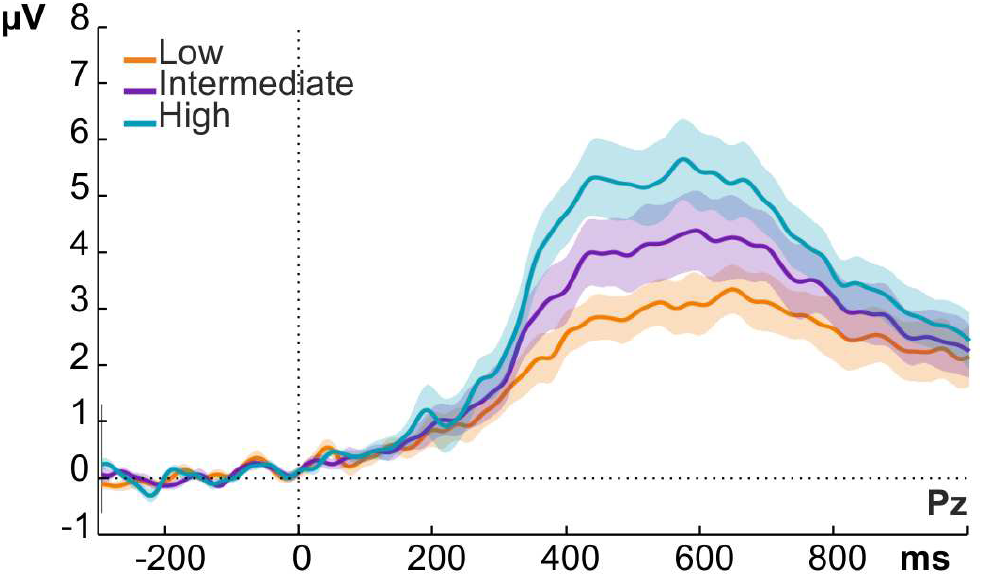
Grand average ERP waveforms over electrode Pz showing the modulation of late evoked potentials by stimulus intensity, regardless of perceptual rating. Shaded areas represent standard errors at each time point

### Centro-parietal positivity co-varies with subjective clarity when stimulus contrast is equated across levels of comparisons

We compared CPP amplitude between awareness ratings for which we could equate stimulus intensities with a sufficient number of trials using trial subsampling (see “Statistical Analysis” section for details). PAS=0, PAS=1 and PAS=2 were compared for stimuli presented at low and intermediate stimulus intensities (Fig. 4a) and PAS=1, PAS=2 and PAS=3 for stimuli presented at intermediate and high stimulus intensities (Fig. 4b). The results were identical for both comparisons (cf. Fig. 4a vs. b). A late positive deflection was observed over central electrodes, peaking around 400-600 ms, which varied with subjective awareness ratings (Fig. 4a-b), increasing in amplitude with stronger subjective experience. To test whether the late positive deflection significantly scaled with awareness ratings, we ran a non-parametric cluster-based permutation test ([17, 18], see methods). This revealed a positive cluster over centro-parietal electrodes between roughly 200-800 ms after stimulus onset (highlighted by the dashed rectangle in Fig. 4a-b), independently of the awareness levels compared (see Fig. 4c-d, left maps for the results of the initial, random sub-sample, p_cluster_ < 0.01). We corroborated this result by repeating the analysis in another 500 runs, randomly selecting a different subset of trials on each iteration (and always equating stimulus intensity across awareness ratings), revealing this effect to be highly consistent across sub-samples (see Fig. 4c-d, right maps for the topography of averaged p-values across the total of 500 runs). Additional, pairwise post-hoc comparisons between each awareness level, performed through cluster-based permutation t-tests averaged over the significant time window identified in the main analysis, showed that the CPP differed in amplitude across all awareness levels (Fig. 4e-f). Overall, these results show that the CPP is strongly related to subjective clarity of the percept, with higher amplitudes corresponding to higher clarity of perceptual experience.

**Figure 4.**
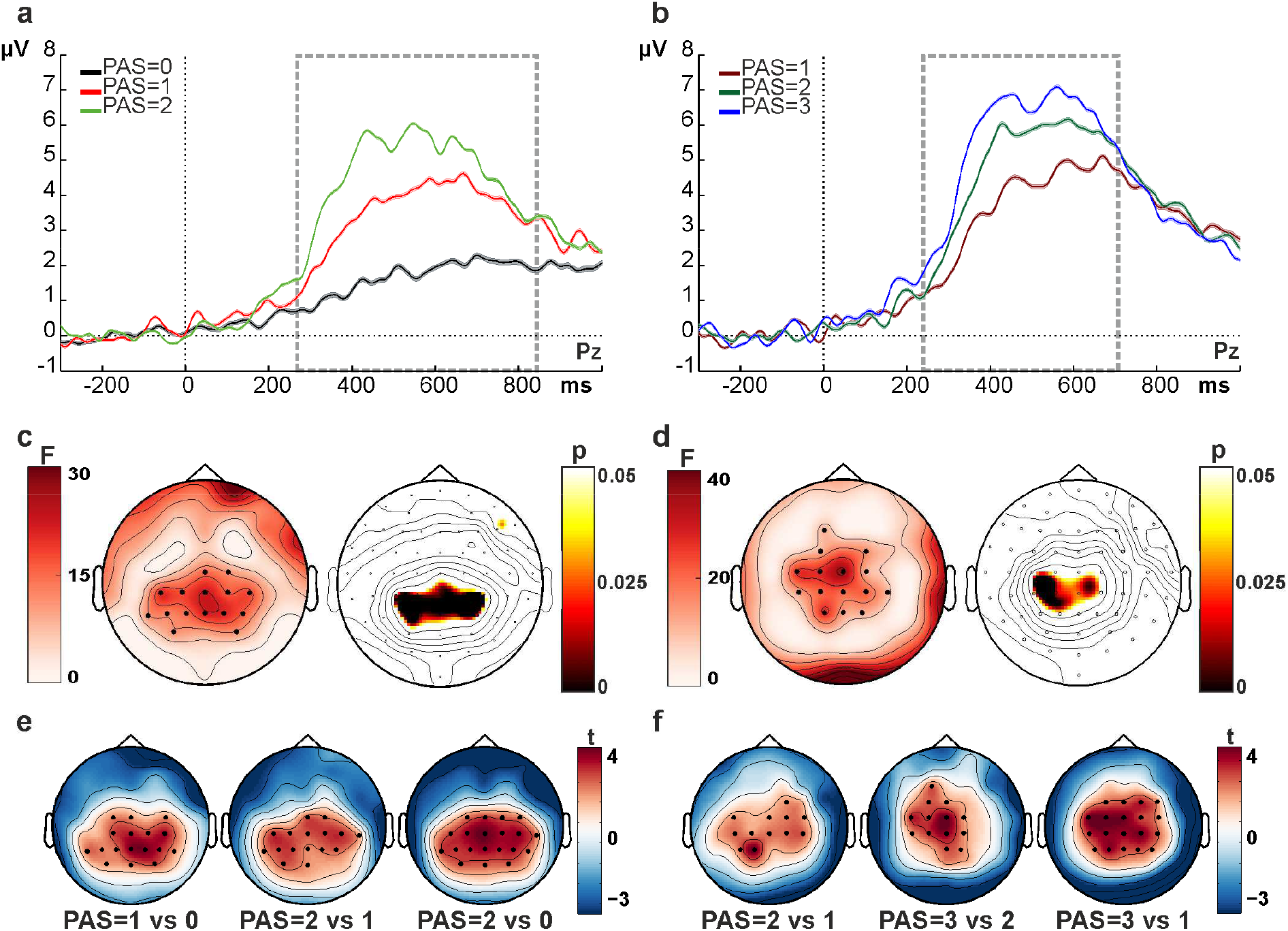
CPP scales with the strength of subjective evidence. **(a-b)** Grand average ERP waveforms over electrode Pz, obtained for each visual awareness rating (**a**: PAS=0 vs. 1 vs. 2; **b**: PAS=1 vs. 2 vs. 3) for trials with equated stimulus intensity levels. Each waveform represents the average ERP over 500 random trial draws; shaded areas represent standard errors at each time point (note that the standard errors are very small and therefore almost invisible). Time windows of significant differences are highlighted by the dashed rectangle. **(c-d)** Cluster analysis results for awareness-related signals (based on ANOVAs across awareness levels). The maps on the left show the topographic distribution of F-values, while the black dots represent significant electrodes (initial draw out of 500 subsamples). The maps on the right show the topography of the p-values averaged over all 500 random draws in the time windows where the most consistent effects were found. **(e-f)** Pairwise post-hoc comparisons for awareness-related signals in the significant time window (dashed rectangle in a-b). Each map shows the t-value distribution with black dots indicating significant electrodes.

### Centro-parietal positivity shows no relation to stimulus contrast when subjective awareness ratings are equated across levels of comparisons

Next, we compared CPP amplitude across stimulus intensity levels for which we could equate the value of awareness ratings with a sufficient number of trials through random trial subsampling. Low versus intermediate stimulus intensity levels were compared after equating PAS values 0, 1 and 2 across stimulus categories (Fig. 5a) and intermediate versus high stimulus intensity levels were contrasted after equating PAS values 1, 2 and 3 across stimulus categories (Fig. 5b). The results reveal very small variations of the CPP with stimulus intensity for both comparisons (see Fig 5a-b: compare low vs. intermediate intensity waveform in a; and intermediate vs. high intensity waveform in b), with the differences being an order of magnitude smaller than those observed for the awareness effects (cf. Fig. 5a-b). Running a non-parametric cluster-based permutation t-test per stimulus intensity comparison did not reveal any significant cluster of electrodes (all ps_cluster_ n.s.). The absence of any effect was not driven by trial selection as confirmed by repeating the trial selection 500 times (random draws). Hence, the CPP was not modulated by the different stimulus intensities when accounting for the subjective experience ratings.

**Figure 5.**
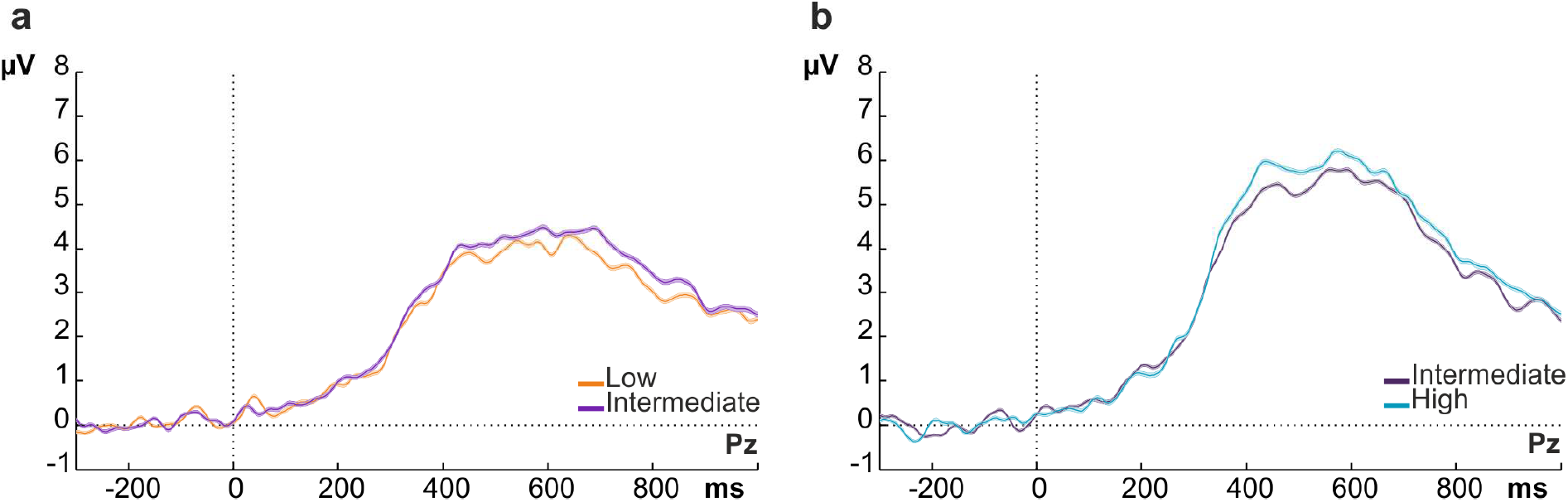
CPP amplitude and stimulus intensity I: ERP amplitudes over electrode Pz per each of the stimulus intensity comparisons (**a**: low vs. intermediate; **b**: intermediate vs. high) for which awareness ratings could be equated. No differences were found across stimulus intensities with equated PAS ratings (compare low vs. intermediate waveform in a, and intermediate vs. high waveform in b). Note that the amplitude-differences between a and b are due to the need for including different PAS ratings in the respective averages (PAS 0, 1 and 2 for waveforms in a, PAS 1, 2 and 3 for waveforms in b). Each waveform represents the average ERP over 500 trial selections (random draws), shaded areas represent standard errors at each time point.

### Centro-parietal positivity: accuracy and catch trials

Given the relatively stronger co-variation of the CPP with subjectively experienced than with physically presented sensory evidence, we expected an uncoupling of this potential from performance accuracy. This was confirmed in two separate analyses. First, we compared the CPP across PAS ratings on correct response trials only (PAS ratings from 0 to 3 included), and then repeated this analysis on incorrect response trials (only including PAS=0 and 1 ratings to have a sufficient number of numerically equated trials between the two categories, see Behavioral results).

For correct trials, we confirmed the existence of an awareness rating effect. The higher the visual awareness rating, the higher the amplitude of the CPP (Fig. 6a, left panel). As above, the cluster-based permutation test with subjective rating (PAS) as a repeated-measures factor, performed on the whole epoch (0 to 900 ms), revealed a significant positive centro-parietal electrode cluster (p_cluster_ < 0.01, from 196 to 900 ms, Fig. 6a, left panel, map). The cluster was also present in each pairwise post-hoc comparison between PAS ratings when performed through cluster-based permutation t-tests on the mean amplitude of the significant time window (p values ranging from 0.046 to 0.002).

**Figure 6.**
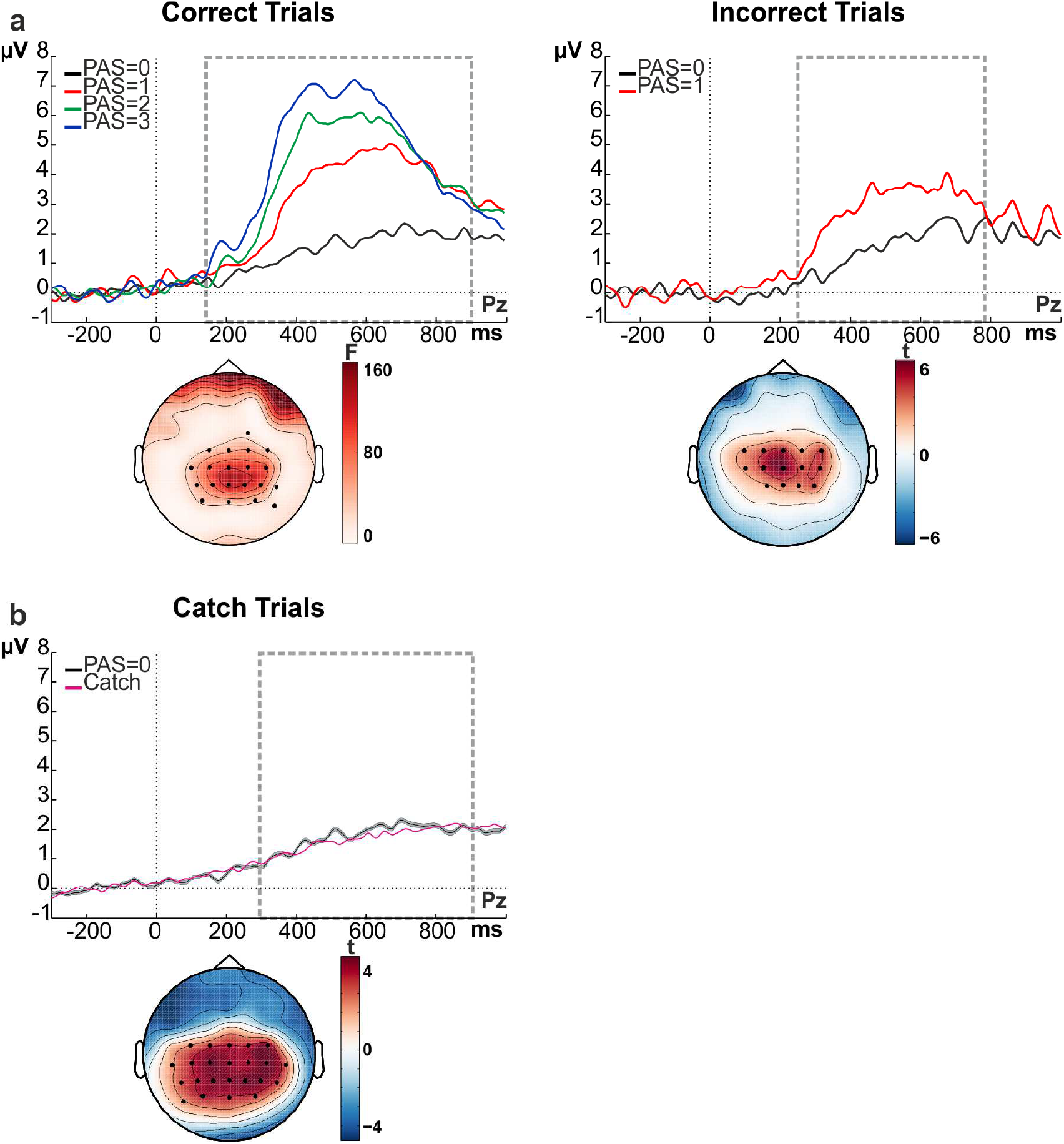
Modulation of late evoked potential by awareness is not related to accuracy. **(a)** ERP waveforms over electrode Pz, obtained for each visual awareness level in correct trials (left panel) and for visual awareness levels PAS=0 and PAS=1 in incorrect trials (right panel). The map illustrates the topographical distribution of F-and p-values for the significant time window (highlighted by the dashed rectangle). Black dots represent significant positive electrode clusters. **(b)** Evoked potentials in catch trials versus PAS=0 trials (obtained after 500 random draws from low and intermediate stimulus intensity trials) over electrode Pz. Shaded areas around PAS=0 waveforms represent standard errors at each time point. Map: t-value distributions from the comparison between catch trial data in the window after expected stimulus onset (marked by 0) versus its baseline “pre-stimulus” interval. Black dots represent significant positive electrode clusters.

Importantly, the awareness rating effect was also observed for incorrect trials, suggesting a dissociation of the effect from task accuracy (Fig. 6a, right panel). The cluster-based permutation t-test (performed on the whole epoch, 0 to 900 ms) again revealed a positive cluster over centro-parietal areas (p_cluster_ < 0.01, from 248 to 788 ms, Fig.6a, right panel, map).

As a variant of testing for a link of the CPP to an ‘internal’ decision quantity, we analysed catch trials to examine whether this late positivity can also occur in the absence of any external sensory evidence. The corresponding cluster-based permutation t-test performed on the whole epoch (0 to 900 ms), with the pre-stimulus period (−300 to 0 ms) as a reference interval, again revealed a positive cluster over centro-parietal areas (p_cluster_ < 0.01, starting at 248 ms until the end of the epoch, Fig. 6b, map). Therefore, even if no veridical sensory information is available, the CPP is still present, and its amplitude matches the amplitude of stimulus-present trials with PAS reports of zero (“no experience”) (Fig. 6b). Please note that the high number of PAS 0 reports in catch trials prevented a comparison across PAS levels.

### Mediation analysis

Based on the initial findings, indicating an effect of the stimulus intensity on PAS report (Fig. 2a) and an effect of the subjective evidence on the CPP amplitude (Fig. 4), we hypothesised that subjective experience may mediate the previously described relationship between stimulus intensity and CPP amplitude. Hence, we tested a mediation model linking stimulus intensity, subjective experience and CPP amplitude [19]. Mediation analysis allows for estimation of the extent to which a proposed mediator variable accounts for the relationship between a predictor and an outcome variable [20]. Here, the stimulus intensity (from catch trials to 75%, 4 levels) was considered as predictor of the EEG amplitude (outcome variable), while the subjective experience measured by PAS scale ratings (0-3, 4 levels) was included as a proposed mediator (see Fig. 7a). In this model, path *a* represents the relationship between stimulus intensity and PAS rating and path *b* the relationship between PAS rating and EEG amplitude when controlling for stimulus intensity. Path *c* represents the total stimulus intensity effect (unmediated) on EEG amplitude and path *c’* represents the direct stimulus intensity–EEG amplitude effect when controlling for PAS ratings. The product of the path *a* and path *b* coefficients (*ab*) represents the mediation effect. Testing for a significant mediation effect involves testing if the predictor-outcome relationship (stimulus intensity – EEG amplitude) is significantly reduced by including the mediator (PAS ratings) in the model (*a*b = c-c’ >0*). To test this, *ab* coefficients calculated for each channel and each time point from 0 to 900 ms after stimulus presentation were tested against 0 by means of a cluster based permutation t-test. We found a significant positive cluster (p_cluster_ = 0.007), spanning from 300 to 800 ms and including several centro-parietal electrodes. Figure 7b shows *ab* values over time averaged across electrodes within the significant cluster (on the left) and its corresponding topography (on the right). Importantly, the time window and the topography of the mediation effect exactly mirrored the CPP component. This result indicates that the inclusion of the mediator “subjective report” in the model significantly decreased the predictive power of the stimulus strength (compare dashed lines in Fig. 7b). In summary, the mediation analysis confirmed and extended our previous results, showing that stimulus strength largely influences the CPP amplitude by affecting the subjective clarity of the percept (Fig. 2a, but also path *a* p < 0.05).

**Figure 7.**
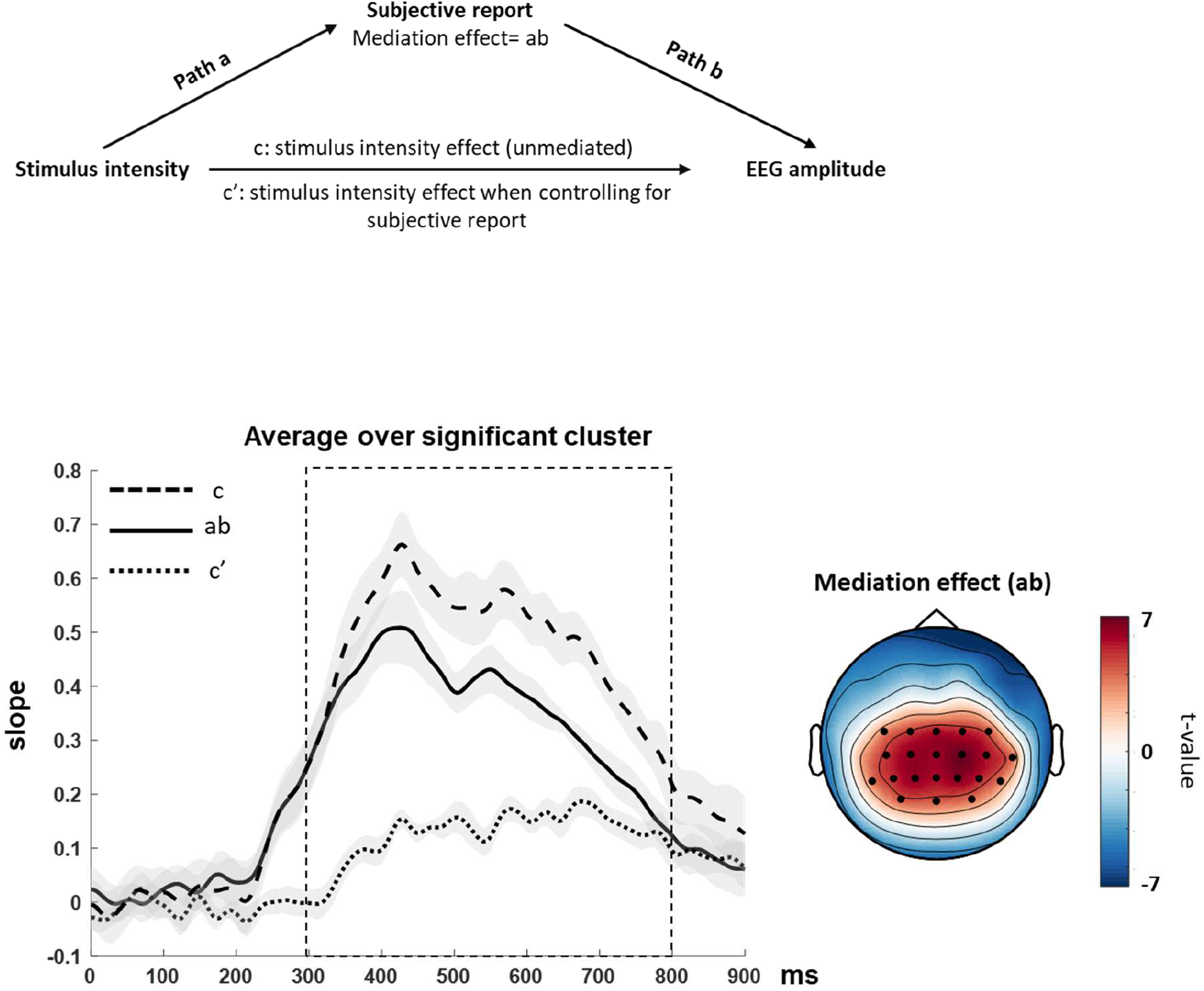
Mediation hypothesis and results. **(a)** The mediation model included the independent variable (stimulus intensity) as predictor of the outcome variable (ERP amplitude). Subjective reports were introduced as a proposed mediator and tested as to whether they accounted for the relationship between predictor and outcome. **(b)** The mediation effect (ab) time course averaged across electrodes within the significant cluster is shown on the left, together with the total effect of the stimulus intensity (c) and the direct effect (c’) when controlling for subjective reports. The rectangle represents the significant time window of mediation. On the right the topography of the mediation effect is shown with the significant positive electrode cluster superimposed (black dots).

## Discussion

Our study sheds light on the relative contribution of objective versus subjective evidence strength to amplitude variations of an EEG evidence accumulation signal (CPP). Our main finding shows a strong preponderance of the latter, internal source to CPP amplitude modulations: when the variation of the CPP amplitude with subjective stimulus clarity is accounted for, there is little residual variation as a function of physical stimulus intensity. In addition, this relationship with subjective evidence strength is observed for error as well as correct trials. Hence, the magnitude of the evidence accumulation signal strongly varies with the subjective experience of stimulus clarity, irrespective of performance accuracy. This provides striking empirical support that the sensory evidence accumulation signal (CPP) closely tracks the individuals’ subjective tally of evidence for their decision.

Our results are highly complementary to a recent study by Kang et al [21], in which participants were asked to report the timing of their commitment to a perceptual decision. By demonstrating that the subjective report corresponds to the time of decision termination, they provided evidence that the neurophysiological mechanisms mediating the decision completion might be also responsible for the piercing of conscious awareness. However, while Kang et al. focused on the timing of the decision process (its termination), we asked our participants to report the clarity of their percept, demonstrating that subjective reports closely match the absolute magnitude reached by the evidence accumulation signal (i.e. the CPP). Interestingly, this suggests that decision processes and mechanisms of conscious perception may be more tightly linked than commonly thought (see also [21]). Moreover, and in further analogy to Kang et al [21], our study suggests a remarkable degree of reliance that can be placed on observers’ subjective impressions to gain access to core internal decision quantities. Intriguingly, a parallel line of research on the neural correlates of consciousness [22] has also identified a late potential of similar topography and latency that tracks conscious reports [23, 24, 25, 26, 27, 28], further substantiating a link between the CPP and conscious perception.

With weak sensory stimulation as employed in the present study, the measure of the clarity of percepts will represent an ‘internal’ subjective quantity, which likely equates to another such quantity: the meta-cognitive measure of confidence in the decision. If participants perceive the stimulus more clearly, they are also likely to be more certain about their decision. A small number of single-unit recordings have demonstrated that neural evidence accumulation signals in parietal neurons index choice confidence, besides representing the choice itself (e.g. [11]). Similar results have been reported in human studies for parietal and frontal sources of activity [29, 12], and for the decision-related P300 component ([30], but see [24] for a possible dissociation of response confidence from awareness). It is unclear whether the awareness rating scale used here relates to the confidence of having correctly detected the stimulus (its presence), or alternatively the confidence in having made the correct choice in the discrimination component of the task (darker/lighter than the background). Given the specific instructions provided to the participants (‘rate the clarity regardless of accuracy confidence’), we tend to favor the former scenario. Precisely how the CPP is related to different types of confidence report (see [31]) is worthy of further study. Irrespective of this, our findings represent a clear demonstration of the link between the CPP and subjective perceptual experience, by relying on the explicit report of subjective awareness, as opposed to indirect behavioral metrics such as post-decisional choice wagering and reaction times [11, 12]. Overall, our findings demonstrate that the CPP reflects an evidence accumulation process, the output of which seems to determine not only the decision itself but also the associated visual awareness levels.

In contrast to a recent series of experiments on the CPP [1, 2, 9, 3], we did not find the late positivity to show a cumulative increase to a common boundary across all conditions, i.e. to display the “build-to-threshold” features of a decision variable. This discrepancy is likely explained by differences in the experimental design. Here and in previous work, the primary task of the participants was to decide which of two stimulus features was presented (e.g. darker vs. lighter dot, left vs. rightward motion patterns, etc.). However, there are important differences in the way the stimuli were displayed, and the responses given. In previous evidence accumulation experiments ([1, 2, 9, 3], see also [32, 33]), sensory stimuli were shown continuously or sequentially (e.g. moving random dot sequences) until a decision was reached, i.e. until all of the evidence needed for a confident report had been ‘accumulated’ and high awareness levels had presumably been reached. In contrast, we used brief static stimuli at peri-threshold levels to capture a range of subjective awareness levels across a range of sensory evidence values. This design allowed us to separate the influence of experienced stimulus clarity from physical stimulus properties.

Our results are in line with previous studies revealing a buildup of the CPP even during erroneous decisions [1, 8, 10] and earlier neuroimaging findings of stronger neural activity for false alarms compared to misses [34] found even at low levels of sensory processing, i.e. in V1-V3 [35]. All of these findings follow from the core concept that random variability in noisy sensory signals impacts on perceptual experience and decisions (see the phenomenon of “choice probability” [36]). Moreover, our data show that the CPP was also present in catch trials, when no visual evidence was physically displayed. The signal recorded during catch trials resembles the shallow evidence accumulation signal observed in monkey parietal areas when no evidence was displayed [37], and replicates previous findings in humans showing a low amplitude late potential in catch trials or when clearly subliminal stimuli were presented [38, 39]. However, it is unclear what this late potential in the absence of sensory evidence reflects, besides that it corroborates that the CPP aligns with an internal quantity. It could reflect an attempt to access evidence for report (i.e. a top-down process), regardless of the presence of the actual physical stimulus, that however fails in most catch trials (around 90%) to reach the threshold for categorizing the percept as a brief glimpse or more (PAS≥1). Interestingly, previous research suggests that a small fraction of catch trials can even be associated with readout of visual information, as indicated by the finding of object perception in pure noise stimuli [40, 41].

### Conclusion

Our results reveal that subjective perceptual experiences closely match the magnitude of the neural signature of sensory evidence accumulation (the CPP) during decision formation. Hence, when probed on clarity of perception, participants may directly ‘read’ the magnitude reached by the decision variable, showing a remarkably tight link between the neural correlates of evidence accumulation and conscious perception.

## 2. Materials and Methods

### 2.1 Participants

14 participants (7 females, 2 left-handed, mean age ± standard deviation: 23.79 ± 3.17) were recruited. All reported normal or corrected-to-normal vision and no history of neurological or psychiatric disorders. They all gave written informed consent to participate in the study, which was approved by the College of Science and Engineering Ethics Committee of the University of Glasgow and conducted in accordance with the 2013 Declaration of Helsinki. Data from three participants were excluded from the analysis because of a low number of trials (< 20) in one or more conditions after artifact removal. The final sample was thus composed of 11 participants (6 females, 1 left-handed, mean age ± standard deviation: 23.6 ± 3.32).

### 2.2 Experimental Procedure

The experiment comprised two sessions performed over two consecutive days. The first session served for calibrating stimulus intensity by threshold assessment (see “Determination of Stimulus Intensity”) and to familiarize participants with the behavioral task. During the second session, after a threshold re-assessment, participants were prepared for EEG recordings. They then performed a forced choice discrimination task, while EEG was continuously recorded (see “EEG Experiment”).

### 2.3 Stimuli

The stimuli were black or white circular patches with a Gaussian envelope (size=1.3°), presented on a gray background in the upper right visual field (5° of vertical and 10° of horizontal eccentricity from the fixation cross). The luminance of the stimuli was individually adjusted to obtain six contrast levels (three lighter and three darker than the background) by means of a threshold assessment procedure (see next paragraph for further details). The contrast of the stimuli presented varied from 0.025 to 0.116% of the maximal contrast of the black/white patch.

### 2.4 Determination of Stimulus Intensity

Participants sat in a dimly lit testing room in front of a CRT monitor (resolution 1280 × 1024, refresh rate of 100 Hz, viewing distance of 57 cm), with their head stabilized by a chin rest. The aim of the first session was to individually determine 3 levels of stimulus intensity (low, intermediate and high) that corresponded to 25%, 50% and 75% of correct detection rate (by manipulating contrast from the medium gray background in both directions, leading to three lighter and three darker patches from background). The thresholds were measured using the method of constant stimuli [42]. At the beginning of threshold assessment, ten evenly spaced contrast values ranging from 0.025 to 0.116% of the maximal contrast of the black/white patches were presented in a randomized order, in the upper periphery of the right visual field (see “Stimuli” for details). This first phase included two blocks: on each block, all contrast values were tested seven times together with 14 stimulus-absent trials (catch trials), resulting in a total number of 308 trials per participant. On each trial, the stimulus appeared after a 1000 ms interval following a brief (150 ms) warning tone (1000 Hz). Participants were asked to keep their eyes on a central fixation cross and to press the spacebar on a keyboard whenever they saw a stimulus. At the end of the two blocks, sigmoid functions were fit to the data from both the light and dark stimulus trials separately and contrast values yielding detection thresholds of 25%, 35%, 50%, 65% and 75% were extracted for each participant. These contrast levels were tested again in two blocks, including 10 trials for each contrast and stimulus type (light and dark stimuli) and 14 catch trials, resulting in a total of 228 trials per participant.

On the second day of testing and prior to EEG recording, a short threshold re-assessment was performed, to verify that participants’ performance was comparable to that obtained in the first session. The previously identified contrast values (5 for light and 5 for dark patches) and contrast levels corresponding to 0% and 100% detection accuracy were each presented seven times together with 14 catch trials, for a total of 182 trials. If contrast values resulting in detection thresholds of about 25%, 50% and 75% were confirmed, they were selected for the behavioral task during the EEG recording. Otherwise, sigmoid functions were again fit to the data and new contrast levels were extracted and tested with the same procedure. The threshold assessment procedure had to be repeated for 4 subjects.

### 2.5 EEG Experiment

During EEG recordings, participants performed a two-alternative forced choice discrimination task. Each trial (Fig. 1) started with a central black fixation cross, followed after 400 ms by a 1000-Hz warning tone (150 ms). After a 1000 ms interval, a light or a dark gray Gaussian patch (the luminance values of which had been determined by threshold assessment) was presented for 30 ms (3 frames) in the upper periphery of the right visual field. A 1000-ms blank screen was then followed by a response prompt asking the participants to judge the brightness of the stimulus relative to the gray background, pressing a button for “lighter” and another button for “darker”. The participants were asked to guess when they did not perceive any stimulus. After the button press, another response prompt asked participants to rate the clarity of their perception on the four-point Perceptual Awareness Scale (PAS; [15]). The four PAS categories are: 0) “no experience” of the stimulus, 1) a “brief glimpse”, 2) an “almost clear experience” and 3) a “clear experience”. Responses were given by pressing four different buttons on the keyboard. The experimental session was divided into ten blocks. Each block was composed of 80 trials: 10 trials for each individually adjusted stimulus contrast (25%, 50% and 75% of detection threshold) and stimulus type (light and dark), together with 20 catch trials, thus yielding a total of 800 trials. The order of the trials was fully randomized. Both the threshold assessment and the actual behavioral task were programmed and run in MATLAB (MathWorks Inc.), using the Psychophysics Toolbox extension [43, 44].

### 2.6 EEG recording and Event-Related Potential (ERP) Analysis

EEG was continuously recorded with a BrainAmp system (Brain Products GmbH, Munich, Germany – BrainVision Recorder) using a Fast’n Easy cap with 61 Ag/AgCl pellet pin electrodes (EasyCap GmbH, Herrsching, Germany) placed according to the 10–05 International System. An additional electrode was positioned below the left eye to record eye movements (after being referenced to Fp1), whereas horizontal eye movements were detected by referencing AF7 to AF8 off-line. Two extra electrodes served as ground (TP9) and on-line reference (AFz). All scalp channels were re-referenced off-line to the average of all electrodes. Electrode impedances were kept below 10 kΩ. The digitization rate was 1000 Hz with a low cut-off of 0.01 Hz and a high cut-off of 100 Hz.

The continuous EEG signal was pre-processed off-line using Brain Vision Analyzer 2.0 (BrainProducts). Data were filtered with a second order high-frequency cutoff of 85 Hz and a second order low-frequency cutoff of 0.1 Hz. A band rejection filter with a bandwidth of 2 Hz was then used to remove 50 Hz interference. Independent component analysis (ICA; [45]) was applied to remove eye blinks and muscle artifacts. The EEG data were then cut into epochs of 1300 ms starting 300 ms before stimulus-onset and baseline corrected using the 300 ms pre-stimulus period. All segments were visually inspected and removed if still contaminated by residual eye movements, blinks, strong muscle activity or excessively noisy EEG. On average, ~5% of the trials were discarded. Finally, data were down-sampled to 250 Hz before averaging.

Analysis of the Event-Related Potentials (ERPs) was performed using the Fieldtrip toolbox ([46]; see http://www.fieldtriptoolbox.org/). Averaging was carried out separately according to visual awareness (subjective rating) and stimulus intensity (detection threshold). To evaluate the unique impact of visual awareness on the late positivity, we randomly selected trials so that each PAS rating would include an equal number of trials with stimuli presented at different detection thresholds, in order to control for the stimulus intensity factor. In a second analysis, we focused on the impact of physical properties of the stimuli, i.e. different intensities. In this case, trials were randomly selected so that ERPs evoked by each stimulus intensity would include an equal number of trials receiving different perceptual ratings on the PAS, to control for the visual awareness factor.

Because of a low number of trials for the low stimulus intensity (25% detection threshold) with rating 3, and high stimulus intensity (75% detection thresholds) with rating 0 combinations, comparisons between perceptual ratings 0, 1 and 2 included trials with stimuli presented at low and intermediate intensity (25% and 50% detection thresholds); comparisons between perceptual ratings 1, 2 and 3 included trials with stimuli presented at intermediate and high intensity (50% and 75% detection thresholds). For the same reason, the comparison between low and intermediate intensity levels only included ratings 0, 1 and 2 and the comparison between intermediate and high intensity levels only included ratings 1, 2 and 3 (Supplementary Fig. 1).

The mean number of trials for each comparison was: 48.55 for rating 0 vs 1 vs 2 and 44.55 for rating 1 vs 2 vs 3; 72.82 for the low vs intermediate stimulus intensities and 66.82 for the intermediate and high intensities.

To investigate the effect of accuracy, correct trials were averaged together according to their awareness rating and regardless of stimulus intensity. For PAS = 0 the mean number of correct trials was 115, 101 for PAS = 1, 112.63 for PAS = 2 and 86.54 for PAS = 3. For incorrect trials, we could only compare PAS = 0 and 1. The difference in trial numbers between the two ratings was overcome by equating the number of trials through a random selection, which resulted in a mean number of trials of 39.81.

Finally, for each participant, average waveforms were computed for the catch trials (mean number of trials after artifact rejection: 186.18).

### 2.7 Statistical Analysis

#### Behavioral data

To evaluate the effectiveness of the experimental manipulations, two separate repeated-measures analyses of variance (ANOVA) were carried out on discrimination accuracy for trials sorted according to either awareness rating (within-subject factor: PAS rating. 4 levels: PAS=0, PAS=1, PAS=2 and PAS=3) or stimulus intensity (within-subject factor: stimulus intensity. 3 levels: low, intermediate and high). We expected higher accuracy for higher ratings and intensity levels.

#### EEG data

To investigate the effect of different levels of visual awareness (PAS=0 vs. 1 vs. 2; PAS=1 vs. 2 vs. 3) and stimulus intensity (low vs. intermediate intensity; intermediate vs. high intensity) on EEG data, non-parametric cluster-based permutation analyses were used [17, 18]. For every sample (channel x time point), conditions were compared by means of a repeated-measures ANOVA (for awareness rating) or by a paired-samples T-Test (for stimulus intensity), on a time window spanning from 0 to 900 ms after stimulus presentation. Those samples whose F-or t-value exceeded a critical value (p < 0.05) were selected and clustered according to spatial and temporal adjacency. Following this, within every cluster, F-or t-values were summed to calculate cluster-level statistics. These cluster-based statistics were evaluated through a non-parametric permutation analysis, which included 500 random sets of permutations. For each permutation, cluster-based statistics were calculated and a reference distribution was built, from which the Monte Carlo p-value was estimated according to the proportion of the randomization null distribution exceeding the maximum cluster statistic. When ANOVAs on visual awareness comparisons were significant, post-hoc analyses were performed through non-parametric cluster-based permutation t-tests between each rating condition. The paired-samples T-Tests were run on the mean amplitude of the significant time window identified by the main ANOVA.

In order to ensure that the random selection of trials performed to equate the number of trials was not biasing the results, trial sampling was repeated 500 times for each comparison and statistical analyses performed for each random draw. The p values obtained after each draw and statistical analysis were averaged together for each comparison, to confirm the significant effects.

To investigate the contribution of stimulus intensity, we compared EEG responses evoked by different amounts of physical information (different stimulus intensities), but resulting in the same subjective report on the PAS. To this end, ERPs evoked by stimuli at low and intermediate intensities and rated as 1 on the PAS were compared to ERPs evoked by stimuli at intermediate and high intensities and again rated as 1. The same comparison was repeated for rating 2. For both comparisons, 500 paired-samples t-tests were run on the mean amplitude of a 900 ms time window (from stimulus onset) on Pz, the electrode that showed the largest effects in the main analysis.

Furthermore, a cluster-based permutation t-test was performed on catch trials to test whether the amplitude of evoked activity significantly differed from a baseline period (−300 to 0 ms before stimulus onset) including all channels and time points from 0 to 900 ms after stimulus presentation.

To investigate the influence of discrimination accuracy on the visual awareness effect, ERP amplitudes evoked by different visual awareness ratings in correct trials only were compared by means of a cluster-based permutation ANOVA (500 permutations, time window from 0 to 900 ms after stimulus onset). Finally, incorrect trials receiving a rating of 0 or 1 were compared by means of a cluster-based permutation t-test (500 permutations, time window from 0 to 900 ms after stimulus onset).

Lastly, a hierarchical (two-level) mediation analysis was carried out using the Mediation Toolbox (http://wagerlab.colorado.edu/tools [19, 47]). First, each path was estimated for each participant, channel and data point (from 0-900 ms after stimulus presentation) using linear regression. The single-trial stimulus contrast values (coded as 0,1,2,3) were entered into the model as the predictor (X) variable and single-trial EEG amplitudes were entered as the outcome variable (Y). Single-trial PAS ratings were tested as a possible mediator (M) of the X-Y relationship. To test for a systematic mediation effect (ab) (i.e. if ab slopes across participants are significantly different from zero), we ran a cluster-based permutation t-test against 0, including all channels and data points.

## ACKNOWLEDGEMENTS

This work was supported by a Wellcome Trust Award to Gregor Thut [grant number 098434]. We would like to thank four anonymous reviewers for their very insightful and constructive comments on a previous version of the manuscript.

## AUTHOR CONTRIBUTIONS

CFT, DV, CSYB, RC, SS and GT designed the research; CFT and DV performed the research; CFT, DV and GT analyzed the data; CFT, DV, CSYB, RC, SS and GT wrote the paper.

---

**Supplementary Figure 1.**
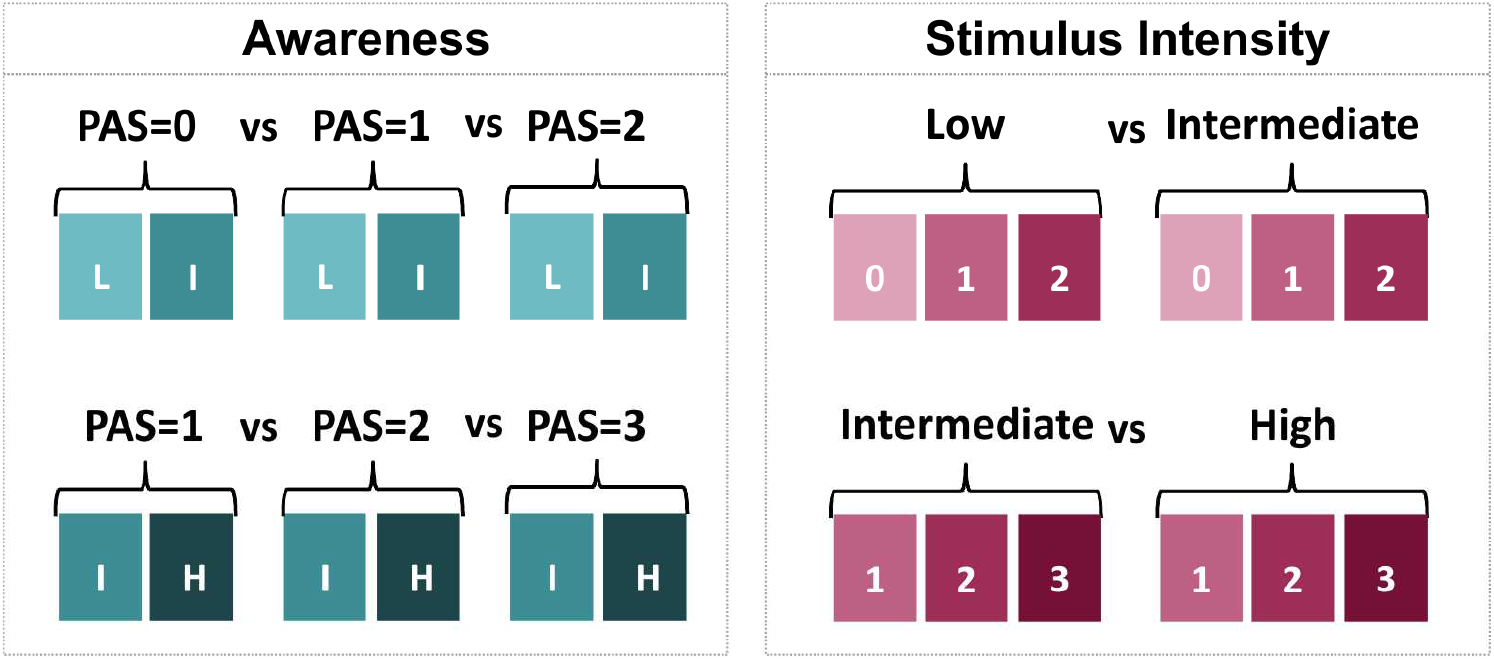
Trial sampling and comparisons. Left panel: comparisons performed to investigate the visual awareness effect. Different rating scores were compared by including an equal number of trials for each stimulus intensity (low (L), intermediate (I) and high (H)). To compare PAS=0 vs. PAS=1 vs. PAS=2, stimuli at low and intermediate intensities were included (left upper panel), whereas stimuli at intermediate and high intensities were included to compare PAS=1 vs. PAS=2 vs. PAS=3 (left bottom panel). Right panel: comparisons performed to investigate the stimulus intensity effect. Different stimulus intensity levels were compared after equating the number of trials for each rating. Stimuli of low and intermediate intensities were compared including an equal number of trials rated with PAS ratings 0, 1 and 2 (right upper panel), whereas intermediate and high intensities were compared including an equal number of trials with PAS ratings 1, 2 and 3 (right bottom panel).

